# Influenza A virus surface proteins are organized to help penetrate host mucus

**DOI:** 10.1101/512467

**Authors:** Michael D. Vahey, Daniel A. Fletcher

## Abstract

Influenza A virus (IAV) enters cells by binding to sialic acid on the cell surface. To accomplish this while avoiding immobilization by sialic acid in host mucus, viruses rely on a balance between the receptor-binding protein hemagglutinin (HA) and the receptor-cleaving protein neuraminidase (NA). Although genetic aspects of this balance are well-characterized, little is known about how the spatial organization of these proteins in the viral envelope may contribute. Using site-specific fluorescent labeling and super-resolution microscopy, we show that HA and NA are asymmetrically distributed on the surface of filamentous viruses, creating an organization of binding and cleaving activities that causes viruses to step consistently away from their NA-rich pole. This Brownian ratchet-like diffusion produces persistent directional mobility that resolves the virus’s conflicting needs to both penetrate mucus and stably attach to the underlying cells, and could contribute to the prevalence of the filamentous phenotype in clinical isolates of IAV.

## Introduction

For a virus to infect a cell, it must reach receptors on the cell surface while avoiding neutralization or clearance by ‘decoy’ receptors in the surrounding environment. For viruses that bind to sialic acid, a receptor that is abundant both on the surface of cells and in the secreted extracellular mucosal environment, attachment and detachment is controlled by receptor-binding and receptor-destroying activities on the viral surface (Rosenthal et al., 1998; Zeng et al., 2008). Influenza A viruses (IAVs), respiratory pathogens that contribute to seasonal flu and have pandemic potential, achieve this balance with hemagglutinin (HA)-mediated receptor binding and neuraminidase (NA)-catalyzed receptor destruction (Air and Laver, 1989; Skehel and Wiley, 2000). Although the importance and genetic basis of the functional balance between HA and NA in IAV has been thoroughly characterized (Wagner et al., 2002; Xu et al., 2012; Yen et al., 2011), it is becoming clear that transmissibility of influenza viruses within and between hosts depends on factors beyond the sequence of these two genes (Chou et al., 2011; Herfst et al., 2012; Neumann and Kawaoka, 2015).

Mucosal barriers present the first line of defense against IAV infection. To infect the underlying epithelium, viral particles must first pass through a ~1-10μm thick layer of mucus that is being steadily transported towards the pharynx where it can be swallowed, neutralizing any virus immobilized within it (Bustamante-Marin and Ostrowski, 2017; Wanner et al., 1996; Zanin et al., 2016). Viruses that bind too tightly to sialic acid will pass through the mucus barrier very slowly, and will thus be unable to reach the surface of an airway epithelial cell before mucociliary clearance. In contrast, viruses that bind only very weakly to sialic acid (or which rapidly destroy it through excessive NA activity (Cohen et al., 2013)) will quickly penetrate mucus, but may be unable to stably attach to the surface of the underlying epithelial cells once it is reached. Adaptations that helps the virus to overcome both of these conflicting challenges, such as changes in virus morphology or a particular spatial organization of envelop proteins, could be evolutionarily favored during in vivo replication.

Interestingly, one feature of IAV that tends to diverge after clinical isolates are cultured in a laboratory environment, or animals are infected with laboratory-grown strains, is particle morphology. While clinical isolates of IAV – samples adapted to transmission in a mucosal environment – form filamentous particles with a consistent diameter but widely varying length, laboratory-adapted strains tend to produce more uniform, spherical particles (Badham and Rossman, 2016; Chu et al., 1949; Dadonaite et al., 2016; Seladi-Schulman et al., 2013). Recent evidence from the 2009 pandemic suggests that filamentous morphology, conferred by the virus’s M segment, may play a role in transmission (Campbell et al., 2014; Lakdawala et al., 2011). However, whether or not virus morphology contributes directly to virus transmission – and if so, how – remains unclear. Similarly, although the two major envelope proteins of IAV, HA and NA, have been observed by electron microscopy to cluster non-uniformly on both the viral and pre-viral envelope (Calder et al., 2010; Harris et al., 2006; Leser and Lamb, 2017), whether and how the spatial organization of HA and NA affects virus transmission remains unclear.

Motivated by these observations, we reasoned that virus shape, together with the packaging and organization of HA and NA in the viral membrane, could influence the balance of attachment and detachment in ways that promote efficient virus transmission. To test this idea, we sought to characterize the organization of proteins in filamentous IAV particles while simultaneously observing their engagement with sialic acid – a challenge with electron microscopy due to the destructive nature of this approach. To overcome this challenge, we recently developed strains of influenza A virus that are amenable to fluorescence microscopy through site-specific tags introduced into the viral genome (Vahey and Fletcher, 2018). Here we show that filamentous particles contain asymmetric distributions of HA and NA in their membranes, and that this distinctive organization biased the diffusion of these particles in a persistent direction over distances of several microns. By enhancing the effective diffusion of a viral particle without reducing the stability of its attachment to the viral receptor, this mechanism could enhance virus transmission across mucosal barriers.

## Results

We first sought to characterize the organization and dynamics of proteins in the viral membrane. By labeling HA and NA, along with the viral nucleoprotein, NP, we are able to measure features of virus organization on intact, infectious particles that corroborate and extend previous observations made using electron microscopy (Calder et al., 2010; Chlanda et al., 2015; Harris et al., 2006; Leser and Lamb, 2017). Images of viruses with labeled nucleoprotein (NP, the most abundant protein in vRNP complexes and a proxy for the virus genome) reveal individual foci of NP that localize to one of the virus’s poles, which we use as a fiducial mark for comparing HA and NA localization along the viral envelope. Out of 1136 filamentous viruses longer than 4.5µm, 540 contained an NP focus at one of the viral poles. We find that NA is enriched ~2x at this pole relative to the middle region of the virus and the opposite pole (Fig. 1A). This protein distribution is stable, since photobleached portions of unfixed filamentous particles do not recover fluorescence in either HA or NA over tens of minutes (Fig. 1B). To investigate finer details of protein organization and to determine if NA in fluorescent viruses is clustered, as suggested by electron microscopy (Calder et al., 2010; Harris et al., 2006), we use two-color stochastic optical reconstruction microscopy (STORM) to reconstruct images of HA and NA with resolution ~10X better than the diffraction limit (~30 nm compared to ~300 nm; Fig. 1 – figure supplement 1). At this resolution, we find that the tendency of NA to concentrate at one of the viral poles is pronounced even in particles smaller than 300nm in length (Fig. 1C). We also find that the NA seen at low levels along the length of filamentous viruses without super-resolution imaging is actually organized into small NA clusters that appear to largely exclude HA (Fig. 1D). Collectively, these measurements present a picture of a variegated IAV envelope whose spatial organization is stable and coupled to the presence and location of the viral genome, with 70% of NP-containing viruses having NA biased to the proximal pole.

**Fig. 1:**
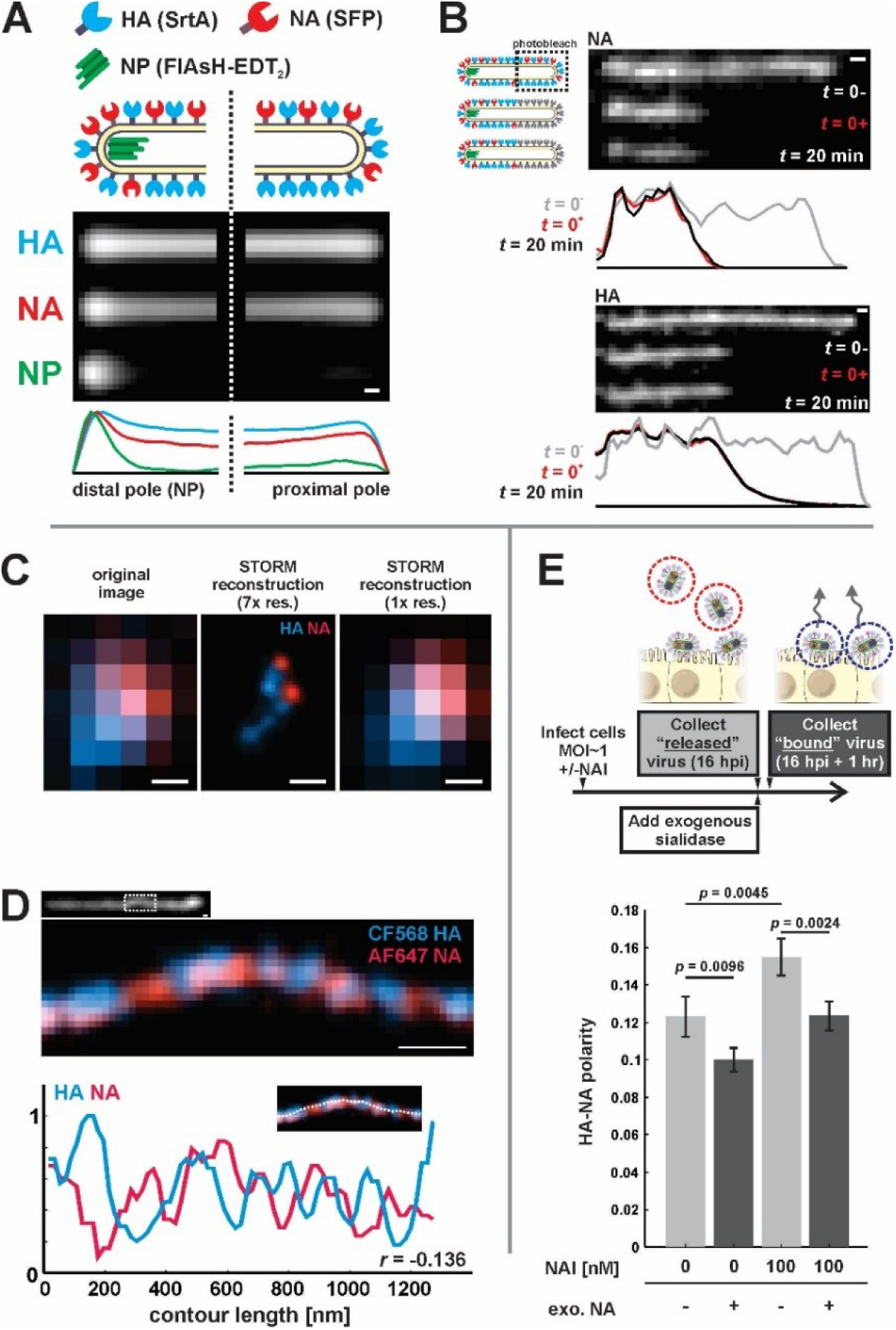
Organization of the IAV envelope in filamentous virus particles. (A) A composite image of the poles of 540 filamentous viruses, aligned by the location of the viral genome (NP). NA intensity at the NP-containing (“distal”) pole of the virus is enriched ~2x relative to rest of the virus after normalizing by HA intensity (scale bar = 200nm). (B) Photobleached fluorescent NA (top) and HA (bottom) on filamentous particles show no recovery after 20 minutes, indicating that NA and HA are immobilized in the viral membrane (scale bar = 500nm). Data is representative of *n* = 8 viruses (NA) and *n* = 5 viruses (HA) from three biological replicates. (C) STORM reconstructions at ~30nm resolution of a pair of viruses unresolveable by diffraction-limited microscopy, each showing the characteristic localization of NA at one end of the virus (scale bar = 200nm). (D) STORM reconstructions of a filamentous virus at ~30nm resolution reveal clusters of HA and NA which partly exclude each other, illustrated by the inverse correlation in HA and NA intensities along the axis of the virus (scale bar = 200nm). (E) To compare populations of virus that are able to detach from infected cell to those that remain bound to the cell surface with or without NAIs, we collect virus in two separate stages: first, we harvest virus from media at 16hpi, followed by the addition of media supplemented with an exogenous, oseltamivir-resistant bacterial neuraminidase. After one hour of treatment with exogenous NA, we harvest virus that had previously remained bound to the cell surface. Virus collected from cells treated with 100nM oseltamivir exhibit significantly more polarized distributions of HA and NA on their surface, while those collected only after treatment with exogenous NA are significantly less polarized. Bars represent median values +/-S.D. for populations measured in four biological replicates with between 499 and 2067 filamentous viruses each (*p*-values determined using a two-sample T-test).

We next investigated whether spatial organization of the IAV envelope could have functional significance for virus binding and detachment. As a first test of this idea, we compared the spatial organization of viruses that were released from the cell surface with those that remained attached after challenging the virus with the neuraminidase inhibitor (NAI) oseltamivir carboxylate (He et al., 1999) (Materials and Methods). Interestingly, viruses that escaped the cell surface under NAI challenge showed significantly higher HA-NA polarity (defined as the separation between the center of masses for HA and NA divided by the virus length) than viruses released in the absence of NAI challenge (Fig. 1E). These results suggest that viruses with NA concentrated at one of the viral poles may be more effective at navigating environments rich in sialic acid.

To directly test how NA polarization might affect virus motion, we characterized interactions between fluorescently-labeled viruses and sialic acid coated coverslips, where the well-defined geometry and density of sialic acid allow straightforward analysis of virus diffusion (Materials and Methods) (Fig. 2A). Because our approach to fluorescently labeling NA preserves its activity (Fig. 2 – figure supplement 1) and viruses harboring fluorophores on both HA and NA preserve ~85% of their infectivity (Vahey and Fletcher, 2018), we could carry out functional assays with the virus and at the same time visualize HA and NA distributions on the viral membrane. Surprisingly, the motion we observed did not resemble randomly-oriented diffusion of the viral particle, but rather persistent, Brownian ratchet-like diffusion, in which filamentous particles with polarized distributions of NA exhibited directed mobility away from their NA-rich pole (Fig. 2B, Supplementary Video 1). Labeling coverslips with Erythrina Cristagalli Lectin (ECL), which binds specifically to the terminal galactose exposed following sialic acid cleavage (Iglesias et al., 1982), revealed the history of virus trajectories and confirmed that virus motion is accompanied by receptor destruction (Fig. 2B). Aligning the trajectories of mobile particles to the orientation of their HA-NA axis reveals that directional mobility can persist for several microns, many times the length of the particle itself (Fig. 2C). By comparing measured trajectories (from Figure 2C) to simulated random walks in which the number of steps and the size of each step matches the observed data but the direction of each step is uncorrelated with previous steps, we estimate that the diffusion coefficient of a polarized virus is enhanced approximately five-fold as a result of directional correlations (Fig. 2D). Consistent with the observation that polarized distributions of HA and NA serve as a determinant for persistent motion and enhanced diffusion, we find that immobile viruses have, on average, less polarized distributions of HA and NA than mobile ones (Fig. 2E).

**Fig. 2:**
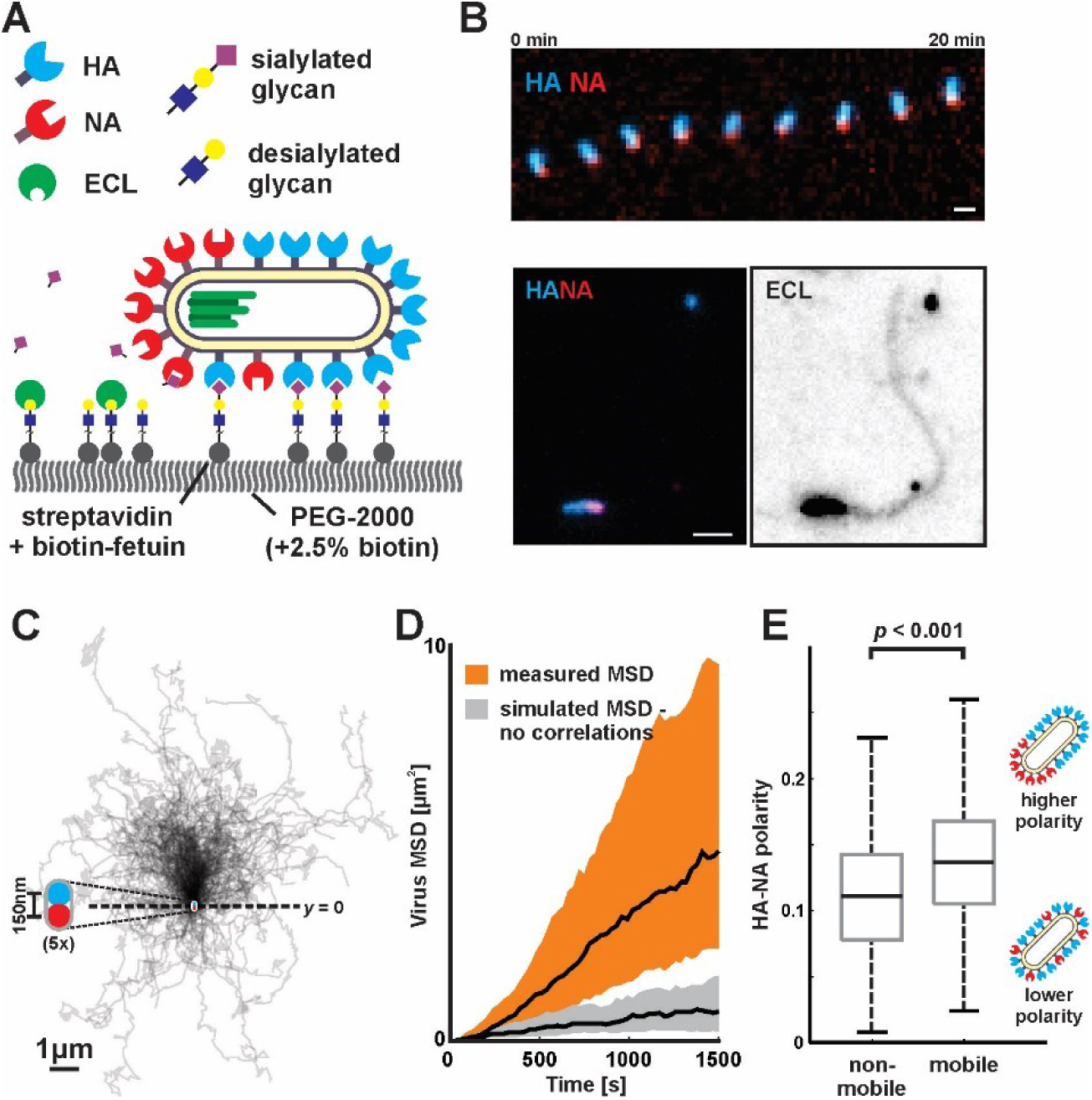
Filamentous IAV diffuses via a Brownian ratchet mechanism. (A) Labeled viruses are placed on coverslips passivated with PEG2K and functionalized with biotinylated fetuin, which provides a high density of receptors for HA and substrates for NA that can be imaged using TIRF microscopy. (B) Time series of a virus migrating in the direction of its higher-HA pole (scale bar = 2µm). Viruses exhibiting persistent motion on sialic acid-coated surfaces leave a trail of terminal galactose to which fluorescent ECL binds, indicating NA cleavage of receptors as the virus moves. (C) Trajectories (measured from timelapse images) of *n* = 240 polarized, mobile viruses, registered to their initial positions and aligned based on the orientation of the HA-NA axis. Blue and red dots at *y* = 0 show the median positions of HA and NA (with median separation of ~150nm), respectively, across all viruses. Data is pooled from three biological replicates. (D) Virus mean squared displacements (MSD) for data from (C), compared to simulated random walks using the same step size and frequency as in (C), but with uncorrelated direction. Black lines show the median MSD, and shaded regions indicate the 25^th^ to 75^th^ percentile range. (E) HA-NA organization correlates with virus mobility. Populations exhibiting little motion (“non-mobile”) have significantly less polarized distributions of HA and NA than those that exhibit persistent directional mobility (quantification of *n* = 403 non-mobile and *n* = 240 mobile viruses combined from three biological replicates; boxes are centered on median values and span from 25^th^ to 75^th^ percentile; *p*-value calculated using a two-sample T-test).

To determine if polarized distributions of NA were necessary for persistent directional mobility, we disrupted the spatial organization of viral surface proteins by removing the cytoplasmic tail of NA (residues 2-6; Fig. 3 – figure supplement 1A) and rescuing a tagged variant of the virus (NAΔCT). Although NA expression on the surface of infected cells is comparable to that seen for wildtype virus, deletion of the cytoplasmic tail reduces packaging of NA into virions (Fig. 3 – figure supplement 1B & C). Additionally, although NAΔCT virus imaged at ~30nm resolution using STORM still exhibits NA clusters resembling those found in wildtype virus, the clusters are no longer immobilized on the surface of the virus, as revealed by photobleaching experiments (Fig. 3 – figure supplement 2A & B). Furthermore, the polarity of NA distributions on the viral surface is significantly decreased in the NAΔCT strain relative to wildtype (Fig. 3 – figure supplement 2C). Using the same sialic acid coated surfaces, NAΔCT viruses no longer exhibited persistent directional motion (Fig. 3A), indicating that spatial organization of NA at the poles is necessary for persistent mobility of IAV. To quantitatively compare the NAΔCT virus to the virus with wildtype NA, we tracked the displacement of viruses between subsequent frames acquired 30s apart and plotted the distribution of the stepping angle relative to the orientation of the virus’s HA-NA axis (Fig. 3B). Viruses with wildtype NA step in the direction of increasing HA (with the NA pole at the rear) roughly twice as frequently as the opposite direction, while NAΔCT viruses exhibit no correlation between orientation and stepping direction (Fig. 3C). Additionally, despite having lower amounts of NA in the viral membrane, NAΔCT viruses left significantly larger trails of cleaved receptors than their wildtype counterparts (Fig. 3 – figure supplement 3). These results demonstrate that the spatial organization and mobility of NA on the viral surface both play an important role in determining where and when it cleaves substrate.

**Fig. 3:**
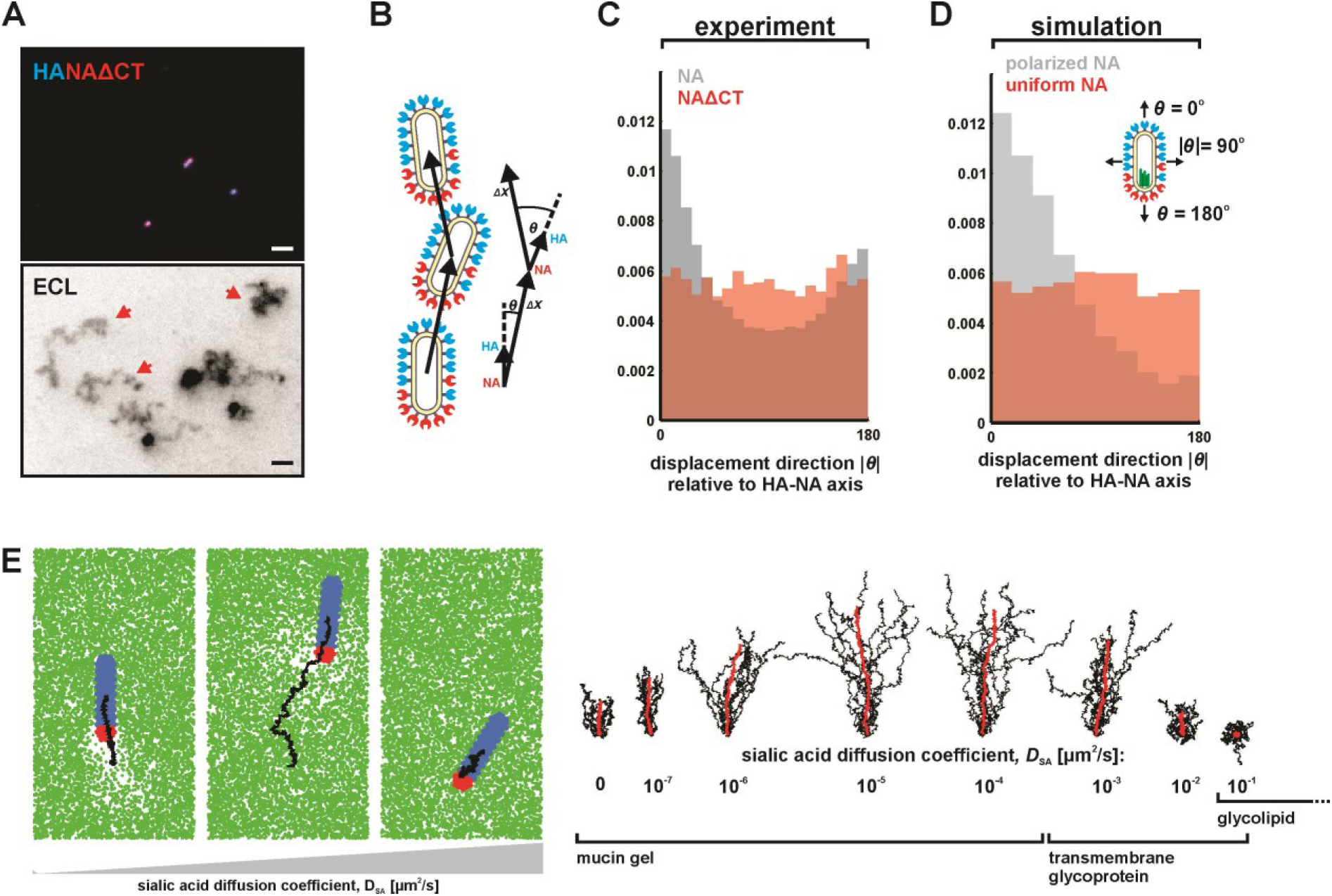
Organization of the IAV envelope and diffusion of the receptor determine persistence of directional mobility. (A) NAΔCT virus create ECL tracks that appear less persistent than those generated by virus with wildtype NA. Red arrows indicate tracks where viruses have dissociated (scale bar = 2µm). (B) Correlations between virus organization and virus motion can be quantified by comparing the angle in which a virus steps relative to the angle of its HA-NA axis. (C) Histogram of virus stepping angle relative to virus orientation for virus with wildtype NA and NAΔCT. While wildtype virus step in the direction of increasing HA roughly twice as frequently as they step in the opposite direction, virus with NAΔCT do not exhibit any orientational preference. Data is pooled from three biological replicates (NA wildtype) or two biological replicates (NAΔCT). (D) Quantification for n = 10 simulations of 250nm viruses with uniform (red) or polarized (grey) distributions of NA, showing the tendency of viruses to step to preferentially along the viral axis only when NA is polarized. The model of an idealized virus is in general agreement with experimental results shown in (C) that include viruses with a wide variety of sizes and HA-NA organizations. (E) The effects of receptor (i.e. sialic acid) diffusion on virus motion. Allowing surface diffusion of sialic acid during simulations can enhance or suppress directional motion of viruses, depending on the receptor mobility and the catalytic rate of NA. Up to a diffusion coefficient of ~0.01µm^2^/s, typical of a slowly diffusing membrane protein, a directional bias in virus diffusion is preserved. Trajectories show the results of 10 simulations for each condition in black, with the average is red.

To investigate the mechanism of this persistent directional mobility, we modeled the diffusion of filamentous viruses with defined spatial organization of HA and NA as they bind to sialic acid on a two-dimensional surface (Supplementary Text, Fig. 3 – figure supplement 4A & B). In this model, virus diffusion is constrained by the tens of attachments the HAs form with sialic acid at any instant in time (English and Hammer, 2004; Xu and Shaw, 2016). At the interface between HA-rich and NA-rich regions of the viral membrane, like those seen in our fluorescence images (Fig. 1A-C), NA will periodically gain access to, and hydrolyze, sialic acid as the virus thermally fluctuates. The binding partners available to HAs located close to this interface will therefore be asymmetrically distributed away from areas of higher NA density. Forming new HA-sialic acid pairs at this interface results in a bound configuration where the virus has more freedom to explore steps normal to the direction of the interface and towards higher sialic acid density than in any other direction. This ultimately results in an increased likelihood of stepping away from the NA-rich pole, which correlates with the overall asymmetric orientation of HA-NA interfaces, and leads to the persistent motion we observe experimentally as well as in our simulations (Supplementary Video 2). Indeed, persistent motion of the virus is lost when NA is no longer localized to the viral poles (Fig. 3 – figure supplement 4C).

Since virus motion is dependent on the virus’s ability to establish distinct regions with and without sialic acid, we expect that dynamic distributions of either NA or sialic acid will alter virus mobility. Consistent with our experimental observations with the NAΔCT virus, allowing NA to freely diffuse eliminates directional bias in virus motion in both experiments (Fig. 3C) and simulations (Fig. 3D). However, allowing surface-bound sialic acid to diffuse can either enhance or suppress directed motion, depending on the kinetic parameters of HA and NA, the diffusion coefficient of sialic acid, and the size of the virus (Supplementary Text). Interestingly, our simulations predict that IAV bound to slowly-diffusing transmembrane glycoproteins (diffusion coefficient <0.01µm^2^/s) or mucins anchored within a gel (diffusion coefficient <0.0001µm^2^/s) will exhibit persistent directed mobility, while virus bound to more rapidly diffusing species (>0.01µm^2^/s) will not (Fig. 3E, Supplementary Video 3). Increasing the length of the virus increases the number of HA-sialic acid interactions, reducing virus mobility but increasing directional persistence (Fig. 3 – figure supplement 5). Thus, the organization of proteins on the viral surface, the size of the virus, and the nature of the receptor to which the virus is bound can each influence the persistence of a virus’s motion.

Although our results focus on a two-dimensional geometry in which a virus adheres to a flat surface decorated with sialic acid, such as a cell surface, we reasoned that they should generalize to three dimensions, such as secreted mucus through which IAV must penetrate to reach naïve cells to infect. To test this prediction, we cultured Calu-3 cells at an air-liquid interface, resulting in a ~1-10µm thick gel of secreted mucus overlaying the apical surface (Fig. 4A). Adding virus with labeled HA and NA to the mucus, followed by fixation and labeling with ECL, revealed tracks similar to those we observed on two-dimensional surfaces, suggesting that the asymmetry of NA and HA biases the direction of virus diffusion in three dimensions (Fig. 4B), producing trails of cleaved sialic acid reaching several microns in length as the virus moves (Fig. 4C). This directional mobility is absent in viruses with more uniformly distributed NA, though they nonetheless create swaths of ECL-staining within the mucus that show no clear directionality (Fig. 4D). These results suggest that the spatial organization of HA and NA on the virus surface may also promote penetration of polarized IAV particles through mucus barriers *in vivo*.

**Fig. 4:**
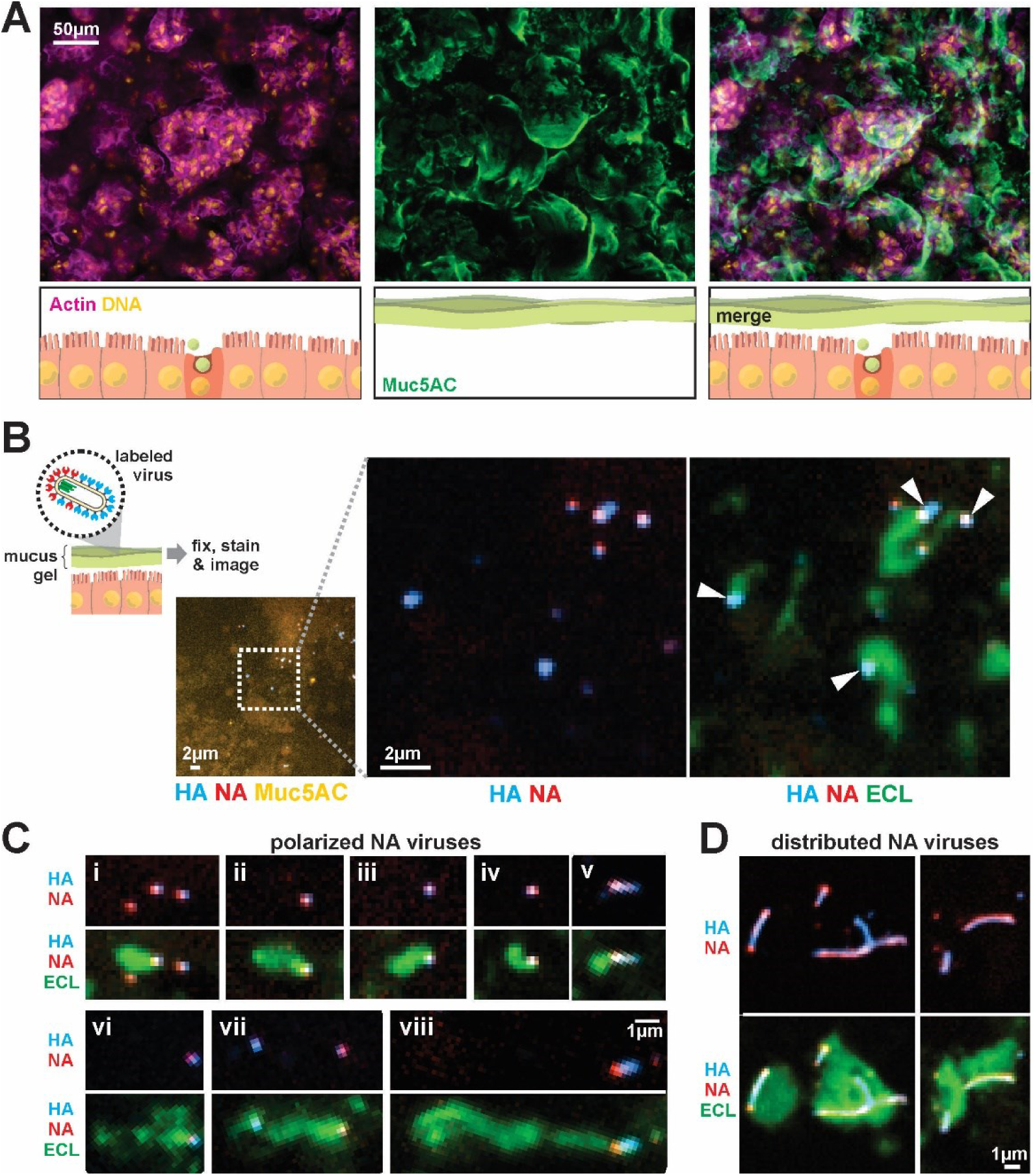
IAV exhibits persistent directional mobility in native mucus gels. (A) CaLu-3 cells cultured at an air-liquid interface for ~10 days partially differentiate, secreted gel forming mucins (visualized with an antibody against Muc5AC). (B) Labeled virus (fluorescent HA and NA) added to the mucosal surface bind and diffuse. Labeling with ECL reveals distinct tracks trailing several microns behind viruses (indicated with white arrows). (C) Panels (i-viii) of registered viruses showing ECL trails. Where polarized distributions of NA on the virus are visible (i, v, vi, viii), ECL labeling is most prominent trailing from the NA-rich pole. (D) Larger filamentous virions with NA broadly distributed across their surfaces clear large patches of sialic acid on the mucosal surface. Images in B, C, and D are a representative sampling from three biological replicates.

## Discussion

Advances in electron microscopy over the past several decades have presented an increasingly detailed picture of the morphology and organization of influenza A virus. This has revealed organizational features of filamentous IAV whose origins and functional significance remain unclear. By using site-specific fluorescent labeling to measure the organization of a virus while preserving its function, we are able to corroborate key observations of envelope protein non-uniformity obtained from electron microscopy and extend them to dynamic observations of both protein motion on the surface of the virus, as well as the surprising finding of persistent influenza A virus motion while engaged with its receptor, sialic acid. These observations demonstrate that the morphology of a virus, the spatial organization of proteins in its membrane, and constraints on the diffusion of these proteins collectively confer the tendency for viral particles to exhibit persistent directional mobility, enhancing the virus’s rate of diffusion without diminishing its strength of adhesion. Importantly, our simulations show that this advantage of filamentous morphology would not be limited to the extremely large particles whose length is easily measured using diffraction limited microscopy, but rather extends to particles <200nm in length (Fig. 3 – figure supplement 5).

Although more work is needed to understand if these characteristics of IAV organization are adaptive during in vivo replication, a simple model of viral transport suggests how the directional viral mobility we observe could contribute during host-to-host transmission. On first entering the respiratory tract, viruses must diffuse through the mucosal barrier to infect the underlying airway epithelial cells. Competing with this process is mucocilliary clearance, which carries particles bound to mucus towards the pharynx where they are neutralized (Fig. 4 – figure supplement 1A). If the transport of a virus through a mucosal barrier is modeled as a first passage problem with a time limit imposed by the rate of mucociliary clearance, a five-fold increase in the diffusion coefficient (as predicted in Figure 2D) would lead to a proportional reduction in the first passage time. More dramatically, it would also increase the fraction of particles that reach the cell surface when the rate of mucociliary clearance is high and would normally prevent most particles from reaching the cell surface. Our experiments on two-dimensional sialic acid-coated surfaces suggest a diffusion coefficient of ~1000 nm^2^/s for polarized viruses; adapting this value for a one-dimensional first passage model suggests that persistent motion increases the number of particles that reach the epithelium before being cleared by ~5000-fold (Supplementary Text, Fig. 4 – figure supplement 1B). Although there are multiple ways that a virus’s passage through mucus can be accelerated (reducing its size, decreasing the number of HAs on its surface, increasing the number of NAs, or making the spatial distribution of NA more uniform), each of these changes would likely reduce binding stability once the virus has managed to reach the cell surface. In contrast, the polarized distributions of NA on the viral surface that we observe increase virus diffusion without compromising binding stability.

Balancing the need to robustly attach to the cell surface with the need to escape immobilization in host mucus is a fundamental challenge for influenza and other viruses that use sialic acid to enter cells. Previous work characterizing the diffusion of influenza C virus, in which receptor-binding and receptor-destroying activities are combined in a single protein (HEF), has shown that the destruction of receptors on a 2D surface prevents virions from retracing their steps (Sakai et al., 2017). Our results show that influenza A virus accomplishes a similar feat through a different mechanism – asymmetric distribution of receptor-binding and receptor-destroying activities on the viral surface that results in persistent motion while maintaining stable attachment. The persistent directional mobility shown here orients filamentous virus motion parallel to its major axis, where it could further enhance the anomalous diffusion coefficients observed for cylindrical nanoparticles in mucus (Yu et al., 2016). Analogous to the combined importance of active motility and cell shape in mucosal bacteria (Bartlett et al., 2017; Sycuro et al., 2010), the directed mobility and cylindrical shape of IAV could help to explain the virus’s ability to penetrate host mucus and contribute to the prevalence of filamentous morphology in clinical isolates of IAV.

## Materials and Methods

### Culturing and labeling virus

Strains of influenza A virus amenable to site specific labeling on HA, NA, and NP were designed and characterized as described previously (Vahey and Fletcher, 2018). Briefly, viruses expressing HA with five consecutive glycine residues following the signal sequence (for labeling via Sortase A (Theile et al., 2013)), NA with a c-terminal ybbR tag (for labeling via SFP synthase(Yin et al., 2006)), and NP with a c-terminal tetracysteine motif (for labeling via direct binding of biarsenical dye FlAsH (Griffin, 1998)) were rescued using reverse genetics (Hoffmann et al., 2000). Viruses used for experiments were collected from confluent monolayers of MDCK-II cells infected at MOI ~ 1 and grown for 16 hours at 37°C in virus growth media (MEM, 0.25% BSA, and penicillin/streptomycin). Media containing virus was collected, centrifuged at 2000×g for five minutes to remove cell debris, and treated with 10mU/ml soluble sialidase (from *C. Perfringens*, Sigma N2876) to ensure that viruses were well dispersed. Viruses were labeled in solution for 90 minutes at room temperature using NTC buffer (100nM NaCl, 20mM Tris pH 7.6, 5mM CaCl_2_) supplemented with 5mM MgCl_2_, Sortase A (180µM enzyme, 50µM CLPETGG peptide) and SFP synthase (5µM enzyme, 5µM CoA probe). Following labeling, Capto Core 700 beads (GE Healthcare; ~1:1 resin volume to sample volume) were used to remove residual dyes and enzymes from the solution of labeled virus. For labeling NP with the biarsenical dye FlAsH, viruses were immobilized on coverslips, washed in NTC buffer, and incubated with 2µM FlAsH for 30 minutes at room temperature.

### Virus photobleaching

To qualitatively evaluate the mobility of HA and NA on the virion surface (Fig. 1B, Figure 3 – figure supplement 2B), unfixed, immobilized viruses were imaged using total internal reflectance (TIRF) microscopy to select longer particles. Filamentous virus of sufficient length (>5μm) were positioned within the field of view such that when the field stop is closed, only approximately half of the virus is exposed to illumination. This half of the virus was then bleached using maximum laser power and then imaged at lower power at 30s intervals as the field stop was opened to monitor recovery. Representative results are shown in Figure 1B and Figure 3 – figure supplement 2B.

### STORM imaging and analysis

Samples for STORM imaging were prepared by binding labeled virus to antibody or sialic acid (i.e. fetuin) functionalized coverslips for one hour at 4°C, followed by the incubation of Dragon Green-labeled 220nm-diameter streptavidin coated beads. Viruses and beads were then fixed to the surfaces with 4% paraformaldehyde in PBS and washed 3x with buffer containing 1M Tris pH 8.0, 5% glucose, and 140mM β-mercaptoethanol. Following these washes, the buffer was supplemented with glucose oxidase and catalase to final concentrations of 0.6mg/ml and 0.035mg/ml, respectively, and mounted on the microscope for imaging.

STORM data was acquired in the following sequence. First, an image of the Dragon Green beads (serving as fiducial marks), HA (labeled with CF568-conjugated peptide and SrtA), and NA (labeled with AF647-CoA and SFP) was acquired, to enable registration. Next, a sequence of STORM images was acquired using a 640nm laser at full power, with acquisition of the HA channel via a 560nm laser at low power every 50 frames to correct for drift. After collecting 15000-35000 frames in this way, we perform STORM imaging on CF568-HA using a 560nm laser. To correct for drift, we acquire an image every 50 frames using a 405nm laser and 575/20nm emission filter; these settings allow us to image the Dragon Green dyes, while simultaneously accelerating blinking of the CF568 dye. For reconstructions of HA, we acquire 25000-35000 frames.

For quantification and localization of blinking events, we use the ImageJ plugin Thunderstorm (Ovesný et al., 2014), combined with custom Matlab scripts for additional drift correction and removal of fluorophores that remain in the “on” state for more than one frame. This analysis results in a list of coordinates for each localization that we then use to reconstruct images of virus at varying resolutions. Reconstructed STORM images (e.g. Fig. 1C & D) are displayed by representing each localization as a gaussian with a standard deviation of 30nm.

### NAI challenge assay

Challenge experiments with the neuraminidase inhibitor oseltamivir are performed as described previously (Vahey and Fletcher, 2018). The analysis from Figure 1E uses an image dataset from ref. 9, reanalyzed to measure the spatial organization of HA and NA on the surface of released virus particles. We infect a polarized monolayer of MDCK cells grown on a collagen gel at MOI of 1-3. After incubating cells with virus for one hour at 37°C, we wash to remove excess virus, replacing media with virus growth media supplemented with or without a specified concentration of oseltamivir carboxylate (Toronto Research Chemicals O700980), but without TPCK-treated trypsin. At 16 hours post infection, we remove the virus containing media for labeling and imaging, and replace with media supplemented with 1U/ml NanI from Clostridium perfringens (Sigma N2876). After treating with this exogenous sialidase for one hour at 37°C, we again collect cell culture media for virus labeling and imaging.

To measure the HA-NA polarization on viruses released in these experiments, we segment filamentous viruses with lengths >1μm and measure the intensity-weighted centroid (i.e. “center of mass”) for both the HA and NA channels. The vector connecting the NA centroid to the HA centroid defines the orientation of HA-NA polarity. Dividing the magnitude of this vector by the total particle length gives the HA-NA polarity metric plotted throughout this work.

### Virus motility assay

Coverslips presenting sialic acid for virus attachment were prepared as described previously (Vahey and Fletcher, 2018). Briefly, NH2-PEG-OH (Rapp Polymere, 122000-2) supplemented with 2.5 mole-percent NH2-PEG-Biotin (Rapp Polymere, 133000-25-20) was conjugated to silanized coverslips for subsequent attachment of sialylated proteins. Following PEGylation, custom PDMS chambers were attached to coverslips, and chambers were incubated for 10 minutes at RT with streptavidin at 50µg/ml in 150 mM NaCl, 25mM HEPEs, pH 7.2, and washed 5X in the same buffer. Fetuin (Sigma F3004) labeled with NHS-biotin was then added at 100nM and incubated ~30 minutes at RT. Coverslips were then washed 5X in NTC buffer and equilibrated to 4°C in preparation for virus binding. For surfaces functionalized in this way (Piehler et al., 2000), we expect a PEG density of ~0.75 molecules/nm^2^, corresponding to roughly one PEG-Biotin per 7nm × 7nm area on the surface. If each biotin is bound by one streptavidin tetramer and subsequently one molecule of biotinylated fetuin (with ~10 sialic acid residues per molecule), the surface density of sialic acid will be ~0.2 SA/nm^2^.

Viruses with HA and NA enzymatically labeled as described previously were bound to coverslips equilibrated to 4oC for one hour on ice. Immediately before imaging, excess virus was washed with pre-chilled NTC, and the sample was mounted on the microscope stage. After allowing the chamber to equilibrate to room temperature, the buffer in the chamber was exchanged to NA buffer (100mM NaCl, 50mM MES pH 6.5, 5mM CaCl_2_), and timelapse recordings were collected using TIRF microscopy at 30 second intervals. To visualize trails of cleaved sialic acid, fluorescein-labeled Erythrina Christagalli lectin (ECL; Vector Labs FL-1141) was added to each well at a concentration of 5µg/ml in NTC buffer with 10mg/ml BSA and incubated for 30 minutes at room temperature. During this incubation period, rapid multivalent binding of ECL to cleaved sialic acid on the viral surface effectively blocks further motion of the virus. Images of ECL trails were acquired using TIRF microscopy without washing unbound ECL from the chamber.

To analyze images of viruses, we separately segment each image according to the intensity in both the HA and NA channels. Merging the two sets of segmented images allows us to determine the position of each virus (i.e. the centroid of the combined HA and NA masks) as well as their morphological features. To determine the HA-NA polarization of each virus, we calculate the distance between the centroid of HA and NA intensity for a particular virus, divided by that virus’s length. The data in Figure 2 and Figure 3 is compiled by extracting these features for each virus in each frame of a timelapse recording.

To analyze the trails of cleaved sialic acid left by mobile viruses, we segment images of samples labeled with fluorescent ECL according to the intensity of lectin staining. Because the virus itself contains high densities of sialic acid on its surface, we use a bandpass threshold to specifically quantify ECL bound to processed glycans on the coverslip (which produces a signal brighter than the background, but dimmer than the virus). Although viruses dissociate from some of the tracks, those that do not allow us to connect intensity of ECL labeling on the surface to intensities of HA and NA on the virus that generated the track. This data is plotted in Figure 3 – figure supplement 3, with and without normalization to HA intensity.

### Quantification of NA activity using MUNANA

For assays using the fluorogenic neuraminidase substrate MUNANA, 10µl of solution containing labeled virus was diluted into 40µl of NA buffer (100mM NaCl, 50mM MES pH 6.5, 5mM CaCl_2_) and incubated in a test tube at 37°C. At time points of 0, 30, 60, 120, and 180 minutes, aliquots were collected and NA was inactivated by adding sodium carbonate to a final concentration of 100mM, and fluorescence was measured by imaging a fixed volume of sample on a confocal microscope using excitation at 405 nm. The rate of turnover was then determined from the slope of the intensity versus time plot with the signal at 0 minutes subtracted from each timepoint. Because strains with wildtype NA and NAΔCT produce different titers of virus that also differ in their morphology and NA composition, we normalized samples to total NA content (Figure 3 – figure supplement 3A), measured by immobilizing viruses on coverslips, imaging them, and integrating the total NA intensity associated with the two samples. This yields an estimated 6-fold difference in total NA content between virus with wildtype NA and viruses with NAΔCT.

### Virus imaging at air-liquid interface of Calu-3 cell cultures

Calu-3 cells grown on plastic dishes for fewer than 10 passages were split at 80% confluence and seeded onto 6mm transwell supports at 50000 cells per insert. Approximately 3 days after seeding, media from the apical compartment was removed (the “airlift”) and cells were provided with fresh media in the basal compartment every other day until being collected for experiments, 8-12 days following the airlift. Cells cultured in this way differentiated into a secretory phenotype, producing a layer of mucus ~1-10µm thick over the apical surface of the monolayer (Fig. 4A). To bind virus without washing away secreted mucus, 5-10µl of labeled virus was added to the apical side of each transwell insert (enough to just coat the surface), and immediately removed, leaving ~1µl of residual virus-containing solution that is approximately evenly distributed on the surface of the monolayer. The cells were then returned to the incubator for 3-6 hours, allowing virus to bind and diffuse, and allowing some of the residual moisture to dry before the cells are collected and fixed on ice for 20 minutes using 4% paraformaldehyde in PBS supplemented with 1mM CaCl_2_. Following fixation, cells and mucus were labeled with MUC5AC monoclonal antibody (45M1; MA5-12178 ThermoFisher) and Erythrina Christagalli lectin labeled with FITC (5μg/ml; Vector Laboratories, FL-1141). Following labeling, the transwell insert was carefully excised with a razor blade and inverted onto a coverslip for imaging.

### Statistics and replicates

Replicates referenced throughout this paper refer to biological replicates, defined as virus collected from separate infected cell cultures, labeled, and assayed as indicated. No statistical methods were used to predetermine sample size. Image data was excluded from analysis in rare cases if the sample drifted on the microscope stage, or if coverslip preparations showed non-specific virus binding. Statistical tests and the number of replicates used in specific cases are described in figure captions. All statistical tests were performed in Matlab R2017b using the Statistics and Machine Learning Toolbox.

## Acknowledgments

The authors would like to acknowledge members of the Fletcher Lab for feedback and technical consultation. This work was supported by the NIH R01 GM114671 (DAF), the Immunotherapeutics and Vaccine Research Initiative at UC Berkeley (DAF), and the Chan Zuckerberg Biohub (DAF). M.D.V. was funded by a Burroughs Wellcome Fund CASI Fellowship. D.A.F. is a Chan Zuckerberg Biohub investigator.

## Contributions

Conceptualization, M.D.V. and D.A.F.; Methodology, M.D.V. and D.A.F.; Software, M.D.V.; Investigation, M.D.V.; Resources, M.D.V.; Formal Analysis, M.D.V. and D.A.F.; Writing, M.D.V. and D.A.F.; Visualization, M.D.V. and D.A.F.; Funding Acquisition, M.D.V. and D.A.F.; Supervision, M.D.V. and D.A.F.

## Supplementary Text

### Simulations of virus binding and diffusion

#### Goals of the model

We wanted to develop simulations that would provide insight into the mechanisms of the directional mobility of influenza A virus. Because individual viruses are highly variable (differing in shape, composition, and organization) and the dynamics of a cylindrical particle close to a surface are complex, we did not seek to develop a model that quantitatively predicts the rates of virus migration that we observe experimentally. Instead, we focused on predicting the general phenomenon of organization-dependent directional mobility, and how this mobility changes when characteristics of the virus or its receptors (i.e. sialic acid, SA) are altered.

#### Modeling HA-SA binding and NA-SA cleavage

We model sialic acid coated surfaces as a uniform random distribution of SA at a density of 0.02 /nm^2^. Although this value is lower than the estimated density in our experiments as well as on the surface of a cell (~0.1-1 SA/nm^2^)(Rosenberg, 1972), it serves as a reasonable approximation given that HAs do not bind all SA glycoforms with similar affinity, and also that not all SAs on the surface of fetuin may be accessible for binding after the protein is immobilized on the surface. For HA and NA distributions on the surface of the virus, we use a triangular lattice with a spacing of 15nm between neighboring spikes, with 84 HAs and 16 NAs proximal to the surface for a filamentous virus 300 nm long. Although we do not explicitly account for the trimeric / tetrameric nature of HA / NA, this should have relatively little effect on our results since each HA can bind up to three SAs and each NA can cleave multiple SAs. Except in simulations where NA is allowed to diffuse on the viral surface (modeling the behavior of NAΔCT that we observe experimentally, Figure 3 – figure supplement 2B), the locations of HA and NA on simulated viruses are static, translating and rotating with the virus as a rigid object. Although this is consistent with our measurements of protein mobility on the viral surface (Fig. 1B), it does not account for rotation of the virus around its minor axis, which could occur in our experiments.

At each time step in a simulation, we identify HA-SA and NA-SA pairs that are within a radius b of each other; we define these SA as being available for either binding (if close to HA) or cleavage (if close to NA). The radius b, which we take to be 7.5nm, models the flexibility of the carbohydrate and protein backbone to which SA is attached, along with the potential flexibility of HA and NA on the surface of the virus. For HA binding kinetics, we used *k*_*on*_ = 400-1000 M^−1^s^−1^ and *k*_*off*_ = 1s^−1^, based on previous measurements(Sieben et al., 2012; Takemoto et al., 1996). The probabilities of an HA-SA pair forming (*p*_*on*_) or losing (*p*_*off*_) a bond in a time interval Δt are therefore approximated by:

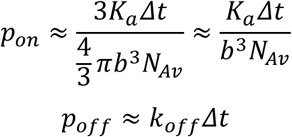

Since the concentration of sialic acid within a binding radius is high relative to the *K*_*m*_ for NA (>10 mM versus 100µM-1mM(Benton et al., 2015)), the probability of an NA-SA pair resulting in cleavage is determined from the catalytic rate constant (*k*_*cat*_ ~ 100s^−1^) as:

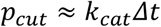

Using these three probabilities, we calculate at each instant in time the number and location of each point of attachment between the virus and the surface and we remove any cleaved sialic acid residues for the remainder of the simulation. Note that we do not prohibit multiple SAs from binding to a single HA centroid; although this would be inaccurate at high densities of SA, under the conditions of these simulations it results in each HA centroid being bound to 0,1, or 2 SAs at any particular instant in time – reasonable values for a trimeric molecule.

#### Modeling virus diffusion

To model diffusion of the virus, we use expressions derived for dilute suspensions of cylindrical rods(Brenner, 1974):

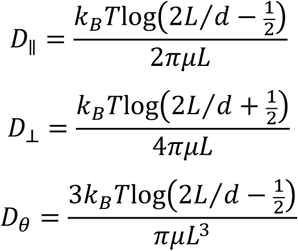

Where *k*_*BT*_is the thermal energy scale, *L* is the virus length, *d* is the virus diameter, and µ is the solvent viscosity. For a diffusing virus constrained to motion in two dimensions with rotation about its long axis but not its short axis, we determine the change in location and orientation during each time interval, Δ*t*, using the following expression(Neild et al., 2010):

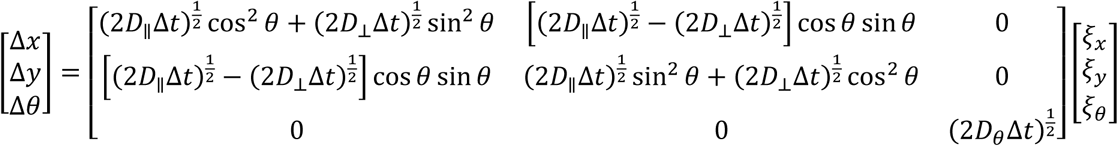

Once the virus has formed attachments to SA on the surface, it’s translational and rotational motion becomes constrained. To account for this constraint, after calculating the translational and rotational increments during a given time interval, we only accept the resulting values if they do not result in extension of any HA-SA bonds beyond their allowable radius, *b*; otherwise, the simulation time does not advance, and we resample the Langevin parameters {ξx, ξy, ξθ} to take another trial step. As the number of bonds increases, the possible steps that satisfy the constraints of all bonds becomes smaller, resulting in immobilization of the virus. In comparison, in the time it would take a single HA-SA pair to dissociate (1/*k*_*off*_ ~ 1s), the typical displacement of an unconstrained virus would be ~1µm, several orders of magnitude larger than would be tolerated by the constraints of the bond. To reduce the number of times we must resample at each time step, we reduce the translational and rotational diffusion coefficients 100-fold to dampen the virus’s motion.

This formulation makes several simplifying approximations, motivated by the cylindrical geometry of viral particles. In particular, we only account for rotations about the virus’ major axis, θ, and not around the angles φ or ψ (Figure 3 – figure supplement 4A). For a cylindrical particle interacting with a flat surface, rotations about the angle ψ would reduce the contact area between the particle and the surface, and would not be favored. This is consistent with our experiments, in which we do not observe filamentous particles pivoting on the surface around the virus’s poles. In contrast, rotations around the angle φ would be permissible, although we are unable to detect them experimentally. Because the particles we model are axisymmetric, rotations about φ do not appreciably change the number or location of HA and NA molecules that are proximal to the surface. As a result, we do not expect that omitting this rotational degree of freedom from our simulations would qualitatively affect the results for cylindrical, axisymmetric particles. However, these simplifying approximations could not be extended to a spherical particle, which could rotate more freely on the surface, altering the number and location of NA molecules capable of cleaving sialic acid.

#### Origin of directed motion

In this model, rectification of virus motion comes from asymmetries in the location of HA relative to the SA that are available for binding. At the boundary between regions where NA activity has cleaved SA and regions where it has not, HA molecules will have more potential binding partners in the direction of the uncleaved region. The constraints introduced by binding to these SAs will leave the virus with greater likelihood to step in the direction of the uncleaved region. Although this asymmetrically distributed freedom of motion will only apply to a subset of all HAs, over the course of ~10^6^ simulation steps, it is sufficient to bias the direction of virus diffusion. This establishes a form of Brownian ratchet, where the input energy could derive from the hydrolysis of sialic acid by NA. Any change in NA organization that gets rid of this asymmetry in the orientation of the HA-NA interface (e.g. placing NA at both poles, in a single band away from the poles, spatially separated from HA, or in a uniform distribution over the surface of the virus) abolishes the rectified motion (Figure 3 – figure supplement 4).

#### Effects of sialic acid diffusion

So far, these simulations have used static distributions of SA. Allowing SA to diffuse on the surface will act to dissipate steep gradients in receptor density created by the activity of NA on the virus, resulting in shallower gradients that extend beyond the localized distributions of NA on the viral surface. The extent of this concentration gradient in available receptors will depend on the balance between the catalytic rate of NA, the rate of diffusion of SA, and the size and rate of motion of the virus. These relationships can be parameterized using a non-dimensional Dahmkohler number:

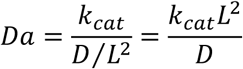

where *k*_*cat*_ is the catalytic rate of NA as before (~100s^−1^), *D* is the diffusion coefficient of sialic acid on the surface, and *L* is a characteristic length for the reaction-diffusion system. Taking *L ~* 100nm, *k*_*cat*_ ~ 100s^−1^, and *D ~* 0.01-0.1µm^2^/s (reasonable values for large glycoproteins(Freeman et al., 2018; Jiang et al., 2015)) gives Da ~ 10-100. The larger this value is, the better able a virus would be to maintain gradients in sialic acid density to support directional mobility. To test this prediction, we performed simulations where the diffusion coefficient of SA is varied while *k*_*cat*_ remains constant and new SA residues are generated at the simulation boundary every time one is cleaved (to prevent complete removal of SA when its diffusion coefficient is high). Consistent with the prediction, simulations of virus motion at Da ~ 100 or greater show preferential diffusion along the axis of the virus, while simulations at Da < 100 do not (Fig. 3E). Interestingly, for IAV this transition corresponds to a diffusion coefficient of ~0.01µm^2^/s, characteristic of some membrane-anchored glycoproteins(Jacobson, 1984), but considerably slower than measured values for gangliosides(Komura et al., 2016). Although receptor diffusion can attenuate persistent virus motion if it is too rapid, it can also increase the rate of motion relative to immobile receptors by allowing HAs away from the HA-NA interface to also sample from asymmetric distributions of receptors (Fig. 3E). Thus, the nature of the receptors to which a virus is bound will likely influence how the virus moves – with greater persistent directional mobility on mucin gels of anchored glycoproteins, and with less persistence once it reaches glycolipids closer to the cell surface.

#### Effects of virus length

Increasing the length of the virus increases the number of potential HA-SA interactions, which we would expect to reduce virus mobility if the kinetics of HA-SA binding are held constant. This trend is supported by simulations; varying particle length (185nm, 285nm, or 385nm) while keeping HA-SA binding kinetics constant dramatically reduces particle mobility (Fig. 3 – figure supplement 5, top row), whereas holding the average number of HA-SA bonds constant results in slightly higher persistent directional mobility for larger particles (Fig. 3 – figure supplement 5, bottom row). In both cases, simulations show a greater tendency for longer viruses to maintain constant orientation. Thus, although particle size and HA-SA binding kinetics both alter virus motion quantitatively, the qualitative features of directional mobility described throughout this work are preserved in particles approaching the lower end of the length distributions observed for filamentous IAV.

### First passage model of virus transport through mucus

The diffusion of viral particles through mucus can be modeled as a one-dimensional problem, where a particle initially binds to the distal part of a mucus gel with thickness Δ and diffuses until reaching the surface of the underlying epithelial cells. Solving the diffusion equation with reflecting boundary conditions at *x* = Δ (modeling the tendency of viruses to stay bound to the mucus layer) and absorbing boundary conditions at *x* = 0 (modeling transfer of the viral particle from the mucus gel to the cell surface) gives the probability density of particles within the mucus gel as a function of time:

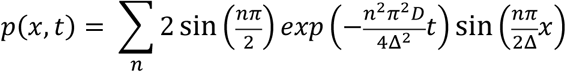

Integrating this expression over *x* gives the probability that a virus will remain in the mucus gel at time *t*, and subtracting this quantity from one gives the probability, *P*(*t*), that a virus will have reached the cell surface by time *t*:

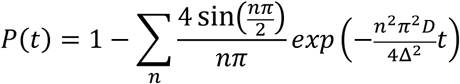

This expression allows us to estimate the survival probability of a virus in the presence of mucociliary clearance by evaluating *P*(*t* = *L*/*U*), where *L* (~10cm) is the distance a bound particle must be carried to be neutralized and *U* (~100μm/s) is the velocity of mucociliary transport. This will depend on the diffusion coefficient, *D*, which we estimate from the data in Figure 2 and the relationship *D* = MSD/4*t*. This gives a value of ~1000nm^2^/s in two dimensions or ~500nm^2^/s in one dimension, which we can substitute into the expressions above. The results of these calculations are plotted in Figure 4 – figure supplement 1, where we compare the first passage probability of viruses with five-fold differences in their diffusion coefficients in a mucus layer that is 2 μm thick. For the parameters listed here, the timescale for mucociliary clearance (~20 min) is substantially shorter than the timescale for virus diffusion (~2h), and the percentage of successful particles is extremely sensitive to their diffusion coefficient: the five-fold decrease in *D* that we estimate would result from the loss of persistent motion would lead to a ~5000-fold decrease in the number of successful particles (Figure 4 – figure supplement 1B). Although this simple model provides only a rough estimate, this analysis, which is based on reasonable estimates of physiological parameters(Bustamante-Marin and Ostrowski, 2017), suggests that persistent motion could enhance the frequency of host-to-host transmission by multiple orders of magnitude.

**Supplementary Video 1.**

Montage of IAV particles acquired using total internal reflectance microscopy at 30 s intervals. Labeled HA is shown in blue and labeled NA is shown in red. Panels are shown at equivalent scales.

**Supplementary Video 2.**

Simulation of a filamentous virus ~250nm in length with a polarized distribution of HA (blue) and NA (red) on its surface bound to a surface coated with sialic acid (green).

**Supplementary Video 3.**

Simulations of polarized filamentous viruses on surfaces with freely-diffusing sialic acid (green). Simulations correspond to *D* = 10^−7^ μ>m^2^/s, *D* = 10^−5^ μm^2^/s, and *D* = 10^−1^ μm^2^/s, as plotted in Figure 3E.

**Figure 1 – figure supplement 1:**
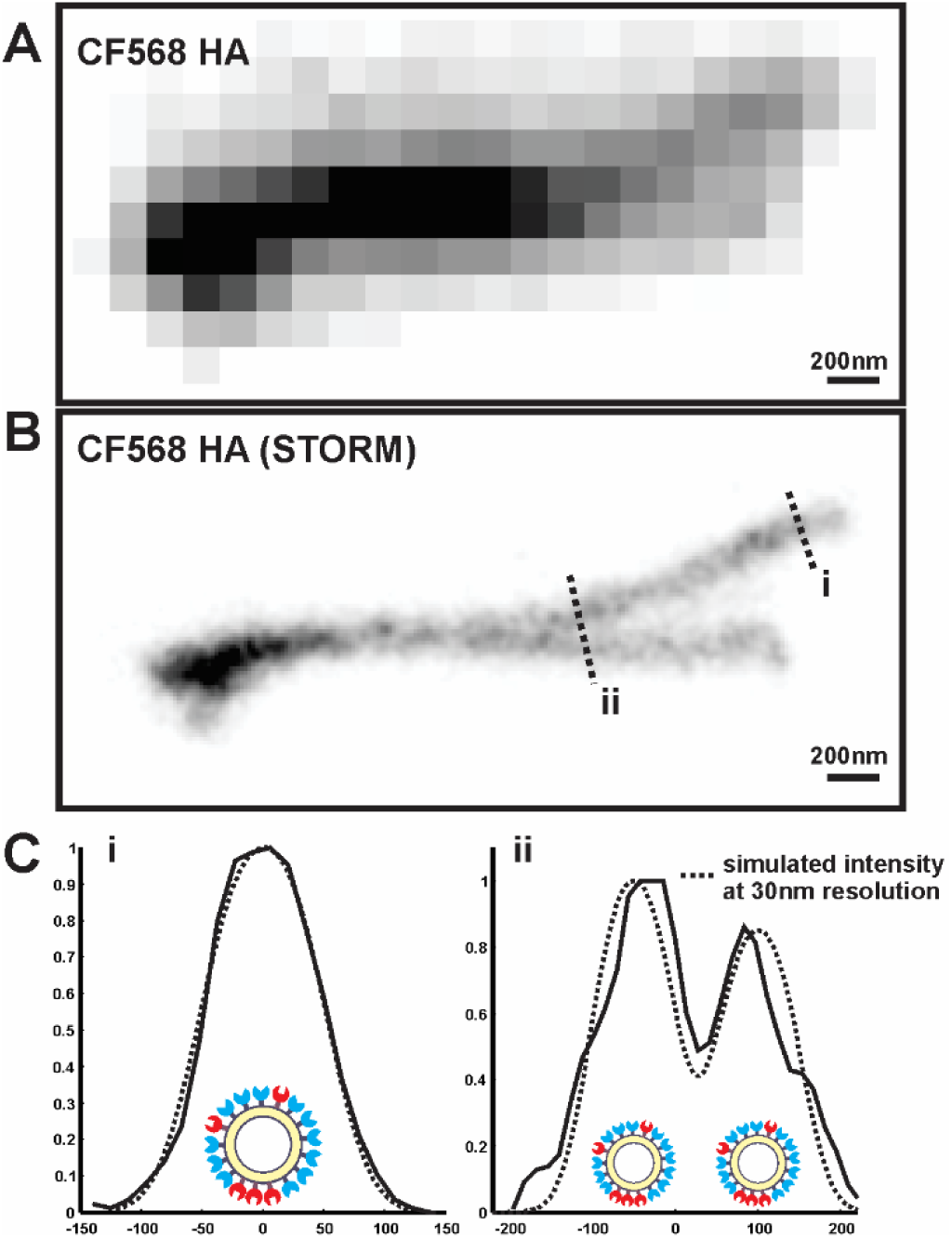
Determining the resolution of virus images reconstructed using STORM. (A) Image of overlapping filamentous viruses acquired using diffraction-limited microscopy (shown with inverted contrast). (B) STORM reconstruction of the same image as in (A), with indicated cross sections I and ii. (C) To estimate the resolution of STORM reconstructions, we compare the collected STORM cross-sections of filamentous viruses (i and ii) to simulated cross-sections, where uniformly distributed molecules on the viral surface are each modeled as a gaussian whose standard deviation serves as a fit parameter. The standard deviation that yields the best fit with our data (~30nm) provides an approximate resolution for our reconstructions.

**Figure 2 – figure supplement 1:**
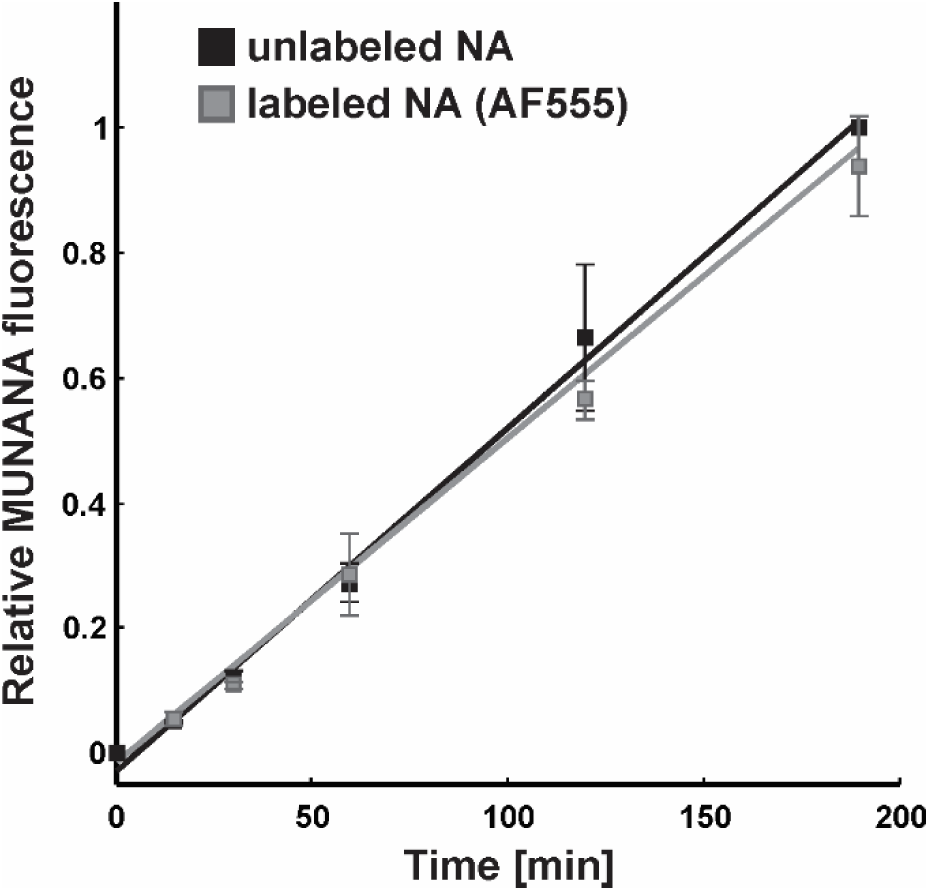
Effect of fluorescent labeling on NA activity. The activity of viruses labeled with Alexa fluor 555 following the same protocol used for imaging experiments matches that of unlabeled samples. Data is from three biological replicates, normalized within each replicate to the MUNANA signal from unlabeled samples at the final timepoint. Under these labeling conditions, the efficiency of fluorophore attachment to NA is >50% (Vahey and Fletcher, 2018).

**Figure 3 – figure supplement 1:**
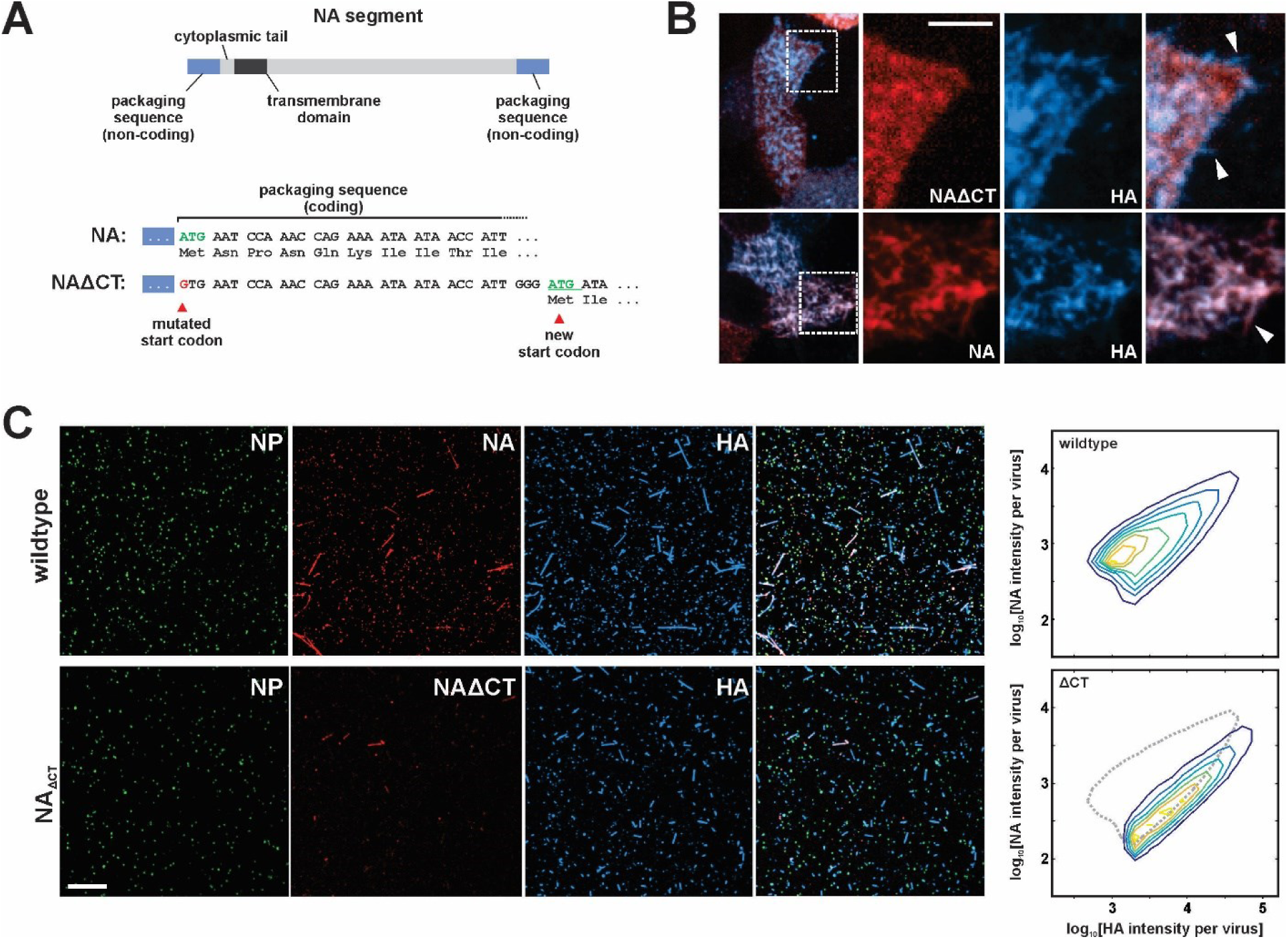
Characterization of influenza A virus lacking the NA cytoplasmic tail (NAΔCT). (A) Construction of an NA segment lacking the N-terminal cytoplasmic tail. Mutating the initial methionine and inserting a new start codon following the coding regions important for vRNA packaging enables the rescue of recombinant virus. (B) NAΔCT virus is expressed on the cell surface but is incorporated into budding viruses (indicated with arrow) less efficiently than wildtype NA (lower panels) (scale bar = 5µm). (C) Imaging released virions supports this observation, with NA incorporation per virion reduced ~three-fold with the deletion of the cytoplasmic tail (scale bar = 10µm).

**Figure 3 – figure supplement 2:**
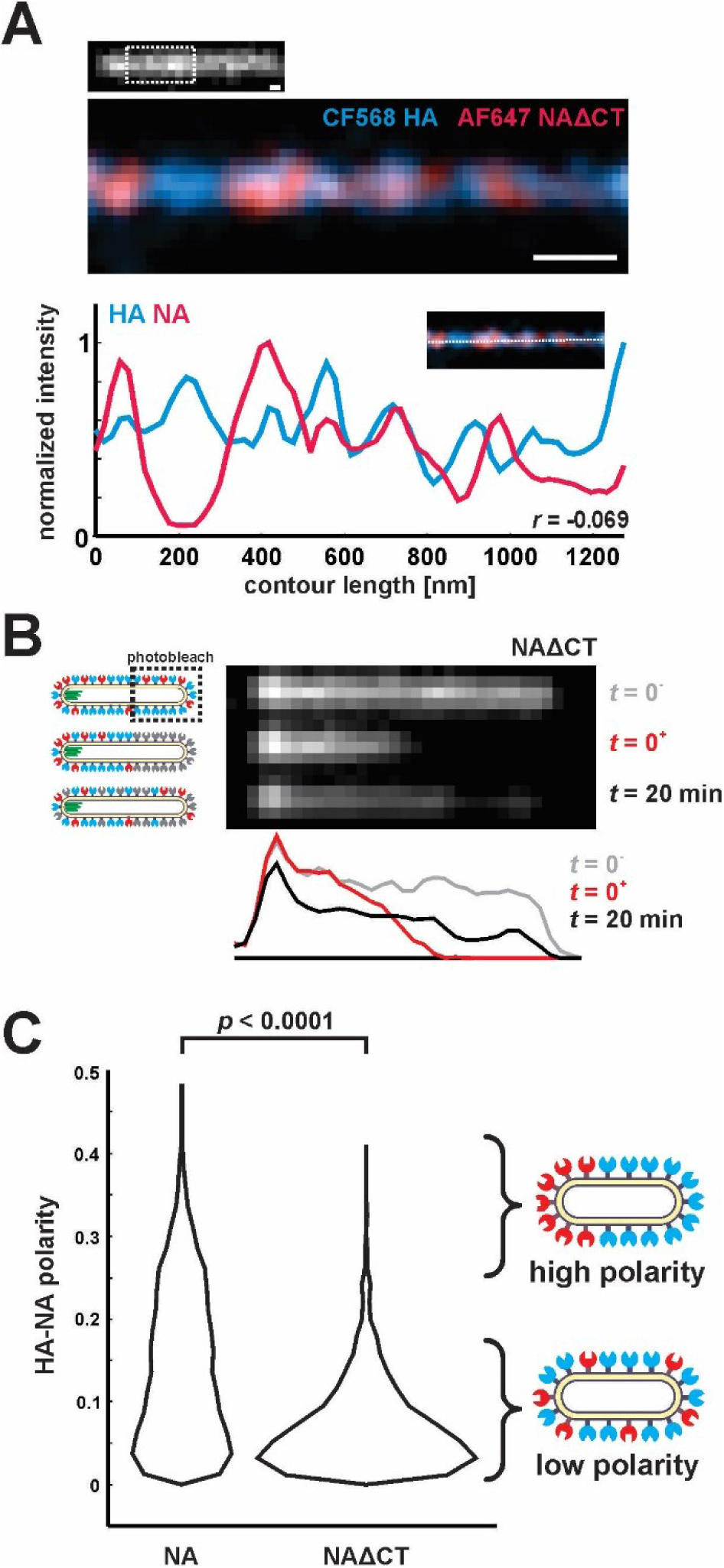
Characterization of NAΔCT virus organization and dynamics in the viral membrane. (A) STORM imaging of NAΔCT reveals a clustered distribution of HA and NA on the surface of filamentous viruses, similar to the nanometer-scale organization of wildtype virus. (B) In contrast to viruses with wildtype NA, photobleaching of NAΔCT on filamentous viruses shows partial recovery on the timescale of several minutes, indicating that the enzyme is at least partially mobile. Results are representative of recovery for n = 8 bleached viruses. (C) Distributions of HA-NA polarity in wildtype and NAΔCT viruses. HA-NA distributions observed in wildtype viruses are significantly more polarized, with NA enriched at the virus poles. Quantification is based on *N* = 6601 wildtype particles (pooled from four biological replicates) and *N* = 2145 ΔCT particles (from one biological replicate); p-value is determined from a two-sample KS test.

**Figure 3 – figure supplement 3:**
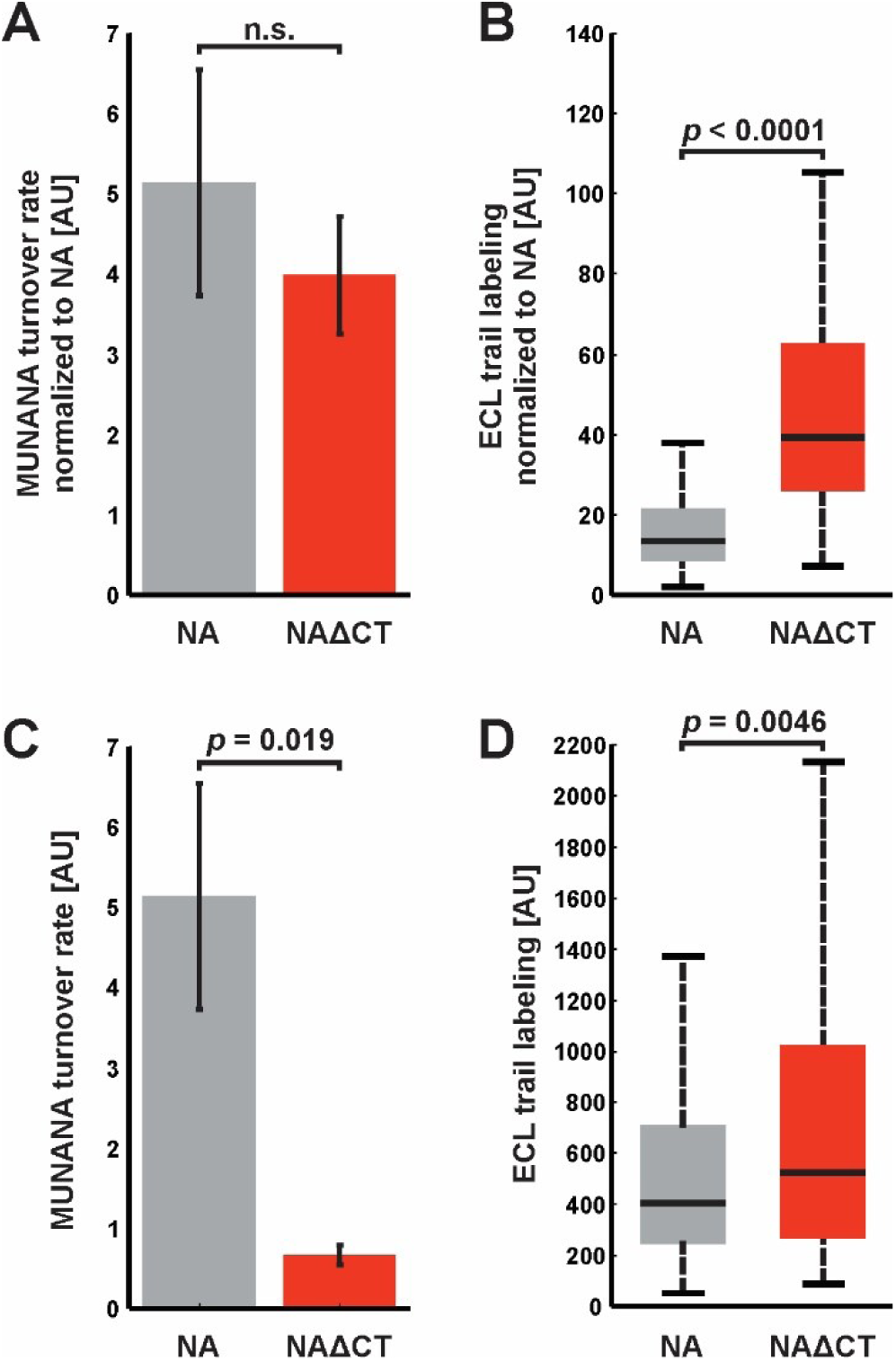
Comparison of NA activities for soluble and immobilized substrates. (A) Enzymatic activity (measured using MUNANA) for intact virus samples with wildtype and NAΔCT, normalized by relative NA content (determined by imaging fluorescently labeled virus). Rates of MUNANA turnover per molecule of NA do not change significantly when the cytoplasmic tail of NA is deleted. Quantification shows the mean and standard error for four biological replicates; p-value determined by a two-sample T-test. (B) Evaluating the NA-activity of individual viruses against immobilized substrates: viruses bound to fetuin-functionalized coverslips were incubated for 30 minutes before labeling with fluorescent ECL to quantify cleavage of sialic acid. Normalizing by NA content (measured by imaging fluorescently-labeled NA) shows a significant increase in the activity of NAΔCT relative to NA(wt) for immobilized substrates. Boxes indicate the median and 25^th^ / 75^th^ percentile measured from two biological replicates with a total of 198 viruses (NA wildtype) and 130 viruses (NAΔCT); p-value determined using a two-sample KS test. (C) and (D) show quantification of the same data, without normalizing for NA abundance for the entire sample (in C) or for individual viruses (in D).

**Figure 3 – figure supplement 4:**
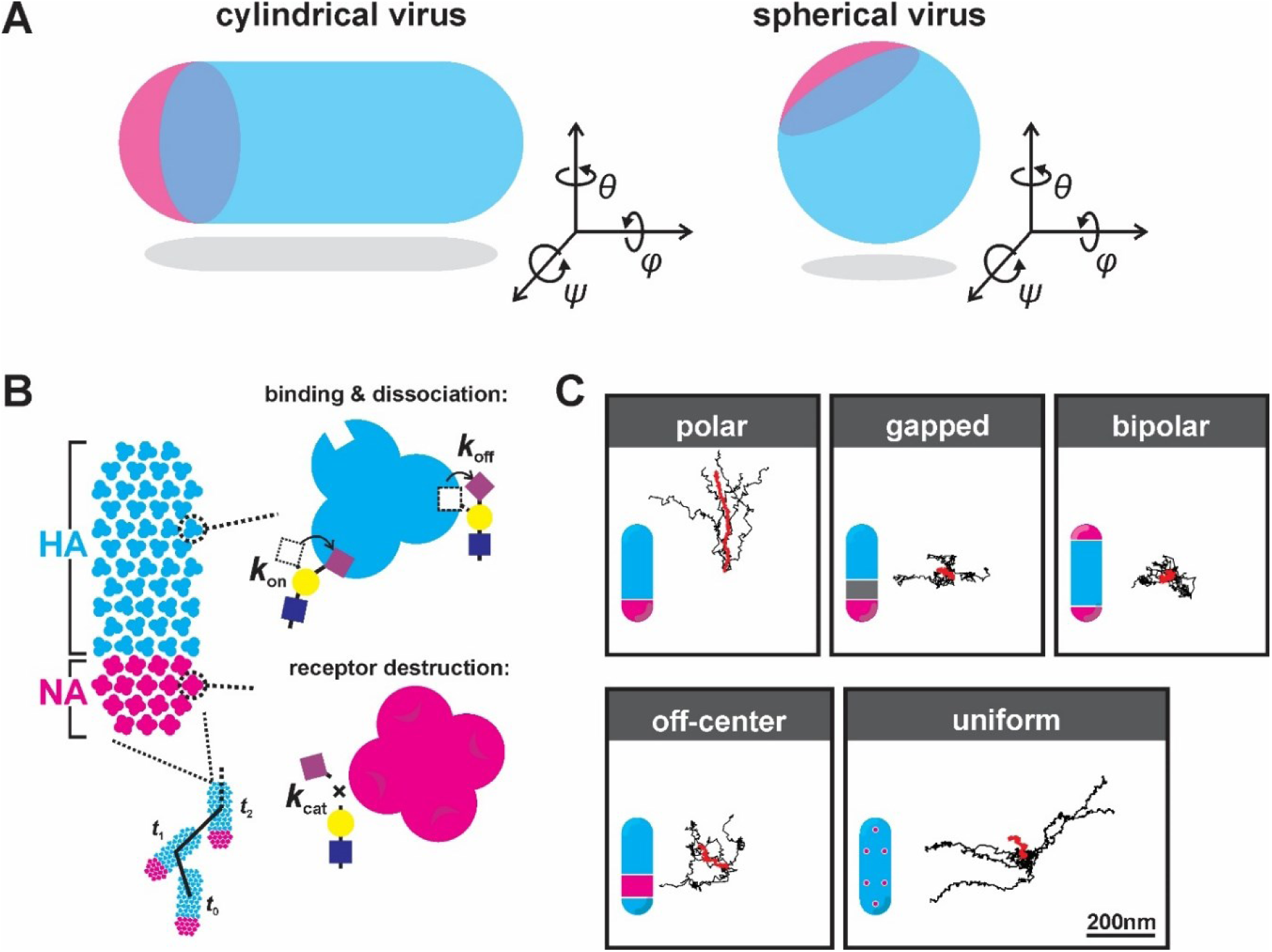
simulating virus binding and diffusion with varying NA organization. (A) Orientation of cylindrical (left) and spherical (right) viral particles on flat surfaces. Of the three rotational degrees of freedom, only rotations in θ are considered in our simulations of cylindrical particles. Rotations in ψ (which reduce the number of possible HA-SA interactions) are not observed experimentally. Rotations in φ likely occur, but do not appreciably change the number and location of HA and NA molecules proximal to the surface. Unlike cylindrical particles, spherical particles would be expected to rotate about θ, φ, and ψ, leading to variations in the number of NAs proximal to the surface over time which would tend to randomize the virus’s direction of motion. As a result, spherical particles are not included in simulations.(B) Model for virus binding and diffusion accounting for multivalent adhesion of HA, turnover of proximal substrates by NA, and thermal fluctuations constrained by the position of bound HA-SA pairs. The virus is modeled as a lattice of HA and NA whose distributions are specified. Viruses interact with uniform distributions of sialic acid on a two-dimensional surface. HA’s within a 7.5nm radius of sialic acid can probabilistically bind, while NA’s within the same radius can probabilistically cleave. Once bound, virus position and bearing can change only if it does not displace any bound HA-SA pairs beyond the specified binding radius (7.5nm in these simulations). (C) Results of simulations with varying organizations of NA on the virus’s surface. Although localizing NA to one of the virus’s poles (‘polar’ configuration) produces consistent oriented motion, introducing separation between HA-rich and NA-rich regions with the same NA localization (‘gapped’ configuration) eliminates this tendency, as does localizing NA to both viral poles (‘bipolar’ configuration). Having moderately polarized NA distributions also suppresses oriented motion, since this configuration (‘off-center’) creates gradients in sialic acid density that partially cancel out over the length of the virus. Finally, NA in a random, uniformly-distributed configuration results in diffusion in a random direction. Because this configuration leads to more efficient cleavage of sialic acid and thus weaker virus adhesion, these simulations were performed with a lower number of total NAs (6 vs. 16) to prevent the virus from detaching altogether.

**Figure 3 - figure supplement 5:**
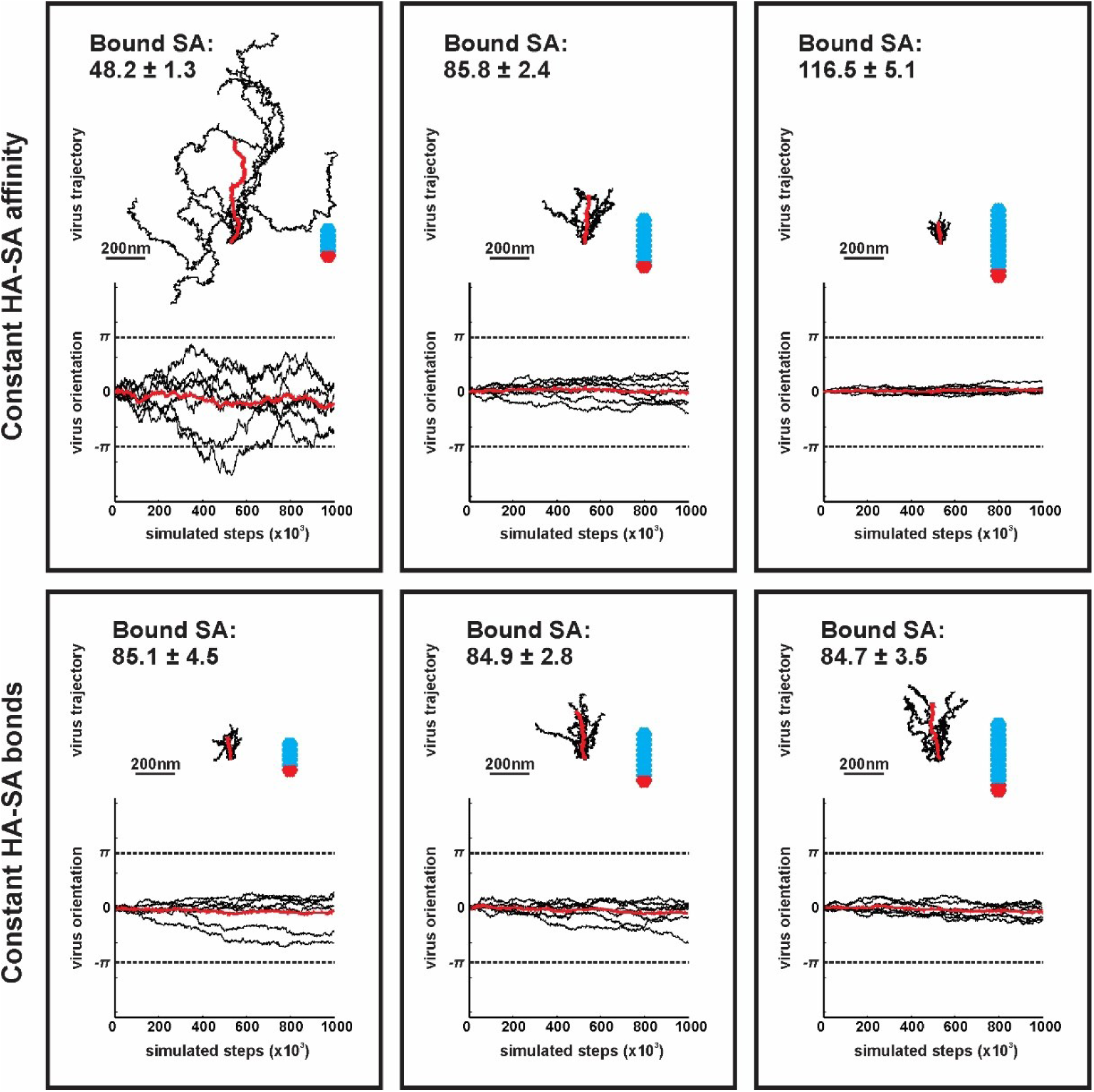
contributions of virus morphology and binding affinity to persistent mobility. Simulations of virus mobility for particles 185nm, 285nm, and 385nm in length, corresponding to 48, 84 and 115 simulated HA trimers per particle, respectively. For a constant HA-SA binding affinity, the average number of bound SA increases proportional to the number of HAs proximal to the surface (top row; average number of bound HA-SA pairs (± S.D.) is given in the upper-right corner of each panel), resulting in substantially reduced mobility in larger particles. If binding affinity is varied (by varying the HA-SA off rate, *k*_*off*_) to maintain a constant number of average bound HA-SA pairs for viruses of different size, larger particles exhibit higher mobilities with greater angular persistence (bottom row). Virus trajectories and orientations are plotted from seven simulations comprised of 10^6^ steps for each of the six conditions shown. Traces plotted in red correspond to averages across all simulations.

**Figure 4 – figure supplement 1:**
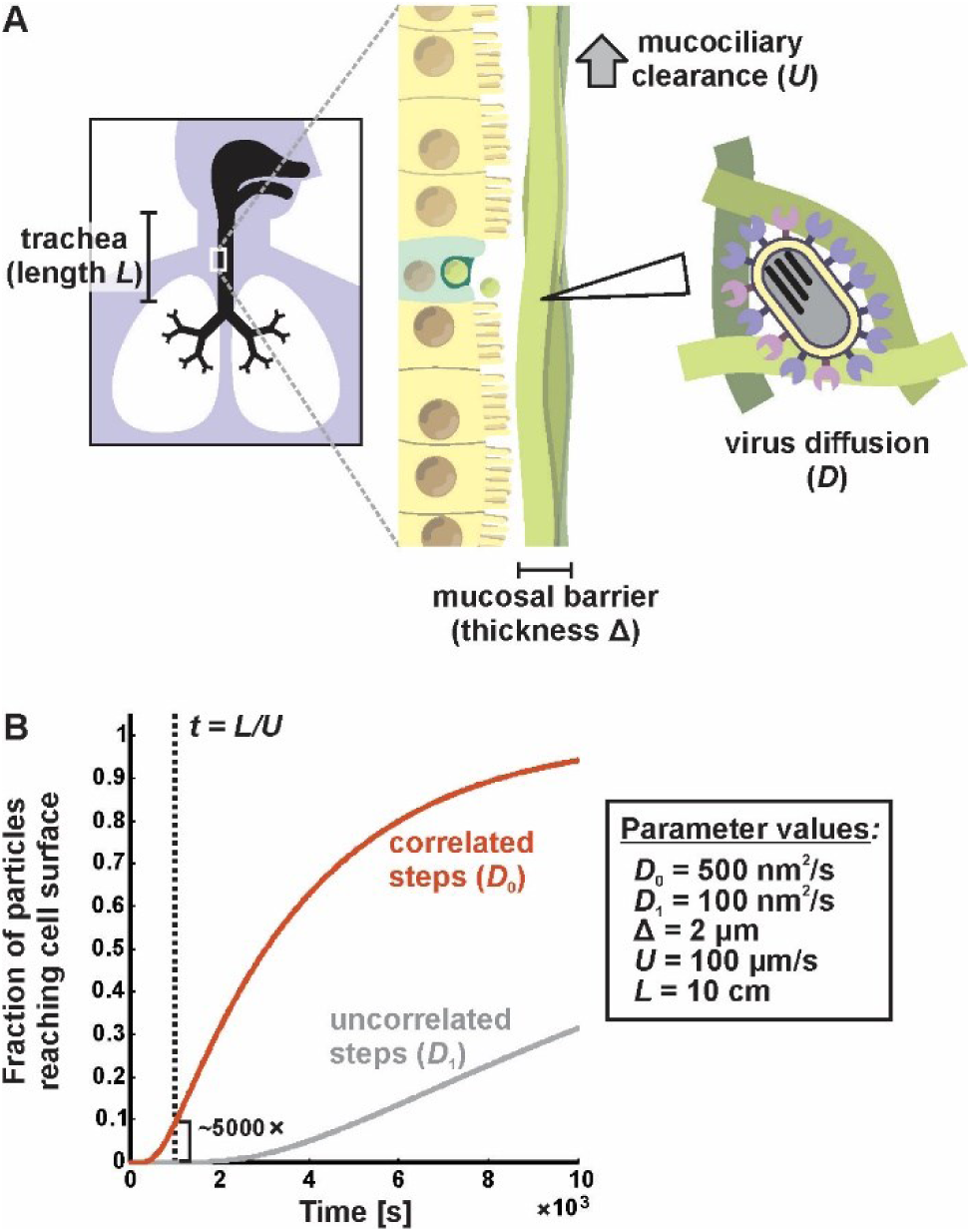
a first passage model for virus transport in mucus. (A) Viruses entering the trachea (e.g. as aerosolized respiratory droplets) bind to the mucosal barrier. The probability of a particle infecting the underlying epithelium depends on the relative rates of virus diffusion (with a characteristic time of ~Δ^2^/*D*) and mucociliary clearance (with a characteristic time of ~*L*/*U*). (B) Solving the first passage problem outlined in (A) for two different diffusion coefficients based on the results from Figure 2 and physiological estimates (inset box) predicts that mucociliary transport is more rapid than virus diffusion, and that a five-fold increase in *D* (as predicted from correlations in the direction of virus motion, Figure 2D) would result in a ~5000-fold increase in the number of particles with an opportunity to infect.

